# High risk of the Fall Armyworm invading into Japan and the Korean Peninsula via overseas migration

**DOI:** 10.1101/662387

**Authors:** Jian Ma, Yun-Ping Wang, Ming-Fei Wu, Bo-Ya Gao, Jie Liu, Gwan-Seok Lee, Akira Otuka, Gao Hu

## Abstract

The fall armyworm, *Spodoptera frugiperda* (J.E. Smith) is an emerging and most severe pest species in the Old World. It is originally native in the Americas. Since 2016 it has spread widely and rapidly to throughout Africa, the Middle East, India, Southeast Asia and most recently southern China. By May 2019 it has appeared in 13 provinces in most southern China and would spread further to northern China. It is highly likely that *S. frugiperda* would enter into Japan and Korea via overseas migrations as many other migratory pests did before. To evaluate the invasion risk of *S. frugiperda* into Japan and Korean Peninsula, we modelled the rate of expansion and future potential migratory range of the insect by a trajectory analytical approach with flight behaviour of *S. frugiperda* implemented, and meteorological data of past five years (2014–2018) used. Our results predicted that *S. frugiperda* would migrate from southern and eastern China into Japan and Korea soon. Most likely, Japan would be invaded from Fujian and Zhejiang on 1 June – 15 July, and Kyushu, Shikoku and south-western Honshu could face the highest risk of *S. frugiperda’*s invasion. Korea would be most possibly reached by *S. frugiperda* from northern Zhejiang, Jiangsu, Anhui, and Shandong on 1 June – 15 July and later. Our results indicated a very high risk that *S. frugiperda* would annually invade Japan and the Korean Peninsula and cause a possible significant decrease in agricultural productivity.

## 1 INTRODUCTION

The fall armyworm, *Spodoptera frugiperda* (J. E. Smith) is an emerging and most severe pest species in the Old World. It is an omnivorous pest native to the tropical and subtropical regions of the Americas (Luginbill,1928; Sparks,1979). *S. frugiperda* has the characteristics of strong migration ability, high reproduction, gluttony, and difficulty in prevention and control (Johnson, 1987; Early, Gonzalez-Moreno, Murphy & Day, 2018; Jiang, Luo, Zhang, Sappington & Hu, 2019). Its host range is extensively wide, including corn, rice, wheat, and more than 180 kinds of other crops, vegetables, ornamental plants, fruits (Casmuz et al., 2010). This insect was known as a super pest of the UN Food and Agriculture Organization Global Alert (FAO, 2018).

With the rapid expansion of international trade, *S. frugiperda* first invaded Nigeria and Ghana in Africa in 2016 (Goergen et al., 2016; Cock et al., 2017). In the following two years, it swept through 44 countries in sub-Saharan Africa (Nagoshi et al., 2018; Rwomushana et al., 2018), caused heavy losses to African food production (Stokstad & Erik, 2017). In May 2018, *S. frugiperda* was first found in India (Sharanabasappa et al., 2018), then spread rapidly to other Asian countries including Thailand, Sri Lanka, Bangladesh, Myanmar, Vietnam, Laos and China (Guo, Zhao, He, Zhang & Wang, 2018; Wu, Jiang & Wu, 2019a; NATESC, 2019ab). On 11 January, 2019, the first confirmation of the invasion of *S. frugiperda* was made in Yunnan Province, China (Jiang & Zhu, 2019; NATESC, 2019a). By May 2019, *S. frugiperda* has spread in most province in southern China, including Yunnan, Guangxi, Guizhou, Guangdong, Hunan, Hainan, Fujian, Zhejiang, Hubei, Sichuan, Jiangxi, Chongqing, and Henan provinces (NATESC, 2019c). Its long-distance migration will significantly increase the risk of spread. According to the forecast results, by July 2019, it is highly possible that *S. frugiperda* would reach the main corn-producing areas of in North China and Northeast China (Wu & Wu, 2019; Li et al.,2019).

The Japanese Islands and the Korean Peninsula as well as eastern China belong to the East Asian migration arena. Their geographical location, ecological environment, and climatic conditions are closely connected each other. Many seasonal pests, such as rice planthoppers (*Nilaparvata lugens, Sogatella furcifera* and *Laodelphax striatellus*) and the oriental armyworm, *Mythimna separata*, can migrate from China to the Japanese Islands and the Korean Peninsula by flying overseas (Otuka, 2013, 2015; Kisimoto & Sogawa, 1995; Hirai, 1995; Lee & Uhm, 1995). The population size of migratory pests in source areas can be directly predict their occurrence level in destination areas (Watanabe, Seino, Kitamura, Hirai, 1990). Now that *S. frugiperda* has arrived in Southeast Asia and southern China, there is a high possibility that they would invade Japan and Korea soon. The invasion of *S. frugiperda* has seriously threatened the production of local corn and other crops and food security. Thus, it is urgent to predict an invasion risk of *S. frugiperda* into Japan and Korea.

To estimate a first-invasion risk of *S. frugiperda* into Japan and Korea, this study conducted a three-dimensional trajectory analysis for recent five years, 2014-2018. We simulated migration paths of *S. frugiperda* to provide the probability of their arrivals. This study presents a scientific basis for monitoring and early warning of *S. frugiperda* in Japan and Korea.

## 2 MATERIALS AND METHODS

We estimated potential endpoints of *S. frugiperda* migrations by calculating forward flight trajectories from source areas where *S. frugiperda* are currently, as of May 2019, known to be breeding, or from potential future source areas we predicted they would reproduce shortly in our previous study (Li et al., 2019). A numerical trajectory model was developed and used successfully for *S. frugiperda* and many other insect migrants, such as the corn earworm *Helicoverpa zea, M. separata*, and the rice leaf roller *Cnaphalocrocis medinalis* (Li et al., 2019, Wu et al., 2018, Wang et al., 2017). This method takes account of flight behaviour and self-powered flight vectors, as these are known to alter trajectory pathways substantially. The trajectory calculation is conducted with high-spatiotemporal-resolution weather data simulated by a numerical weather prediction model.

### 2.1 Weather Research and Forecasting model

The Weather Research and Forecasting Model (WRF) (version 3.8, www.wrf-model.org) were used to produce a high-resolution atmospheric background for trajectory calculation. In this study, the National Centers for Environmental Prediction (NCEP) Final Analysis (FNL) data were 6-hourly, global, 1-degree grid meteorological data that was used as initial field data and boundary conditions for WRF. The hourly initial and boundary conditions were simulated by WRF to run the three-dimensional trajectory program for *S. frugiperda*, which had a spatial resolution of 30 km. Calculation schemes and parameters for WRF modelling are listed in Table 1.

**Table 1.** Selection of scheme and parameters of the WRF model

| Item | Domain 1 |
| --- | --- |
| Location | 32°N, 127°E |
| The number of grid points | 140*150 |
| Distance between grid points(km) | 30 |
| Layers | 30 |
| Map projection | Lambert |
| Microphysics scheme | WSM6 |
| Longwave radiation scheme | RRTMG |
| Shortwave radiation scheme | RRTMG |
| Surface layer scheme | Monin-Obukhov |
| Land/water surface scheme | Noah |
| Planetary boundary layer scheme | YSU |
| Cumulus parameterization | Tiedtke |
| Forecast time | 72 h |

### 2.3 Trajectory modelling of *S. frugiperda*

Based on biological characteristics of the migration of *S. frugiperda*, trajectories were calculated with following parameters. (i) *S. frugiperda* flies downwind at high altitude (Wolf, Westbrook, Raulston & Lingren,1995; Nagoshi, Meagher, Fleischer, 2009), without considering a directional deflection angle (Li et al.,2019). (ii) A self-powered flight vector of 3.0 m/s was added to wind speed in the trajectory modelling (Westbrook et al.,2007; Li et al.,2019). (iii) The noctuid insects fly at night, taking off at dusk, and landing at dawn on the next day (Qi et al., 2013; Wang et al., 2017). According to the sunrise and sunset times at departure points from April to August, take-off time is 20:00 h (CST), landing time is 06:00 h (CST) on the next day; thus *S. frugiperda* flies continuously for 10 h every night, and flies for three consecutive nights as most other noctuid moths, when the moth flies over lands (Wang et al., 2017). (iv) When *S. frugiperda* comes over the sea area, East China Sea and Yellow Sea, during each nocturnal flight, flight duration is extended until it reaches over the land, but does not exceed 36 hours; (v) Because *S. frugiperda* cannot fly when air temperature at flight altitude becomes below 13.8 °C, trajectories that enters into the low temperature zones were stopped on any night/height combination (Hogg, Pitre & Anderson, 1982). (vi) Radar observations show that moth insects usually fly in a low-level jet stream at altitudes with wind speed > 10 m/s (Johnson, 1987; Wolf et al., 1990; Westbrook, Nagoshi, Meagher, Fleischer & Jairam, 2016). In this study, using six different initial heights: 500, 750, 1000, 1250, 1500, and 1750 m above mean sea level (Li et al., 2019), and thus six trajectories were calculated on one night at one departure point.

We simulated the trajectories of *S. frugiperda* by using meteorological conditions at flight altitude from the past 5 years (2014–2018). In total, ∼0.14 million trajectories were calculated, making this the largest study of *S. frugiperda* migration pathways conducted.

### 2.3 Departure points for forward trajectories

Trajectories were started from each potential departure point placed at every 1-degree grid for eight southern and eastern provinces of Guangdong, Fujian, Jiangxi, Zhejiang, Anhui, Jiangsu, Shandong, and Henan. According to the prediction result of *S. frugiperda* in eastern China (Li et al.,2019), it would spread to Guangxi and Fujian in April and emigrate out one month later, and reach Jiangxi, Zhejiang, southern Anhui and southern Jiangsu (south of 32°N) in May and June. At last *S. frugiperda* would cover the entire Anhui, Jiangsu, Henan and Shandong provinces in June and July (Li et al., 2019). Therefore, the eight provinces were divided into following three parts: (i) Guangdong and Fujian from 1 to 31 May; (ii) Fujian, Jiangxi, Zhejiang, southern Anhui (south of 32ºN) and southern Jiangsu (south of 32ºN) from 1 June to 15 July. This time period covers all the dominant rain season in southern China (the *Meiyu* season) and Japan (the *Bai-u* season), and most other insects migrate into Japan during this period (Otuka et al., 2010; Tojo et al., 2013); (iii) Anhui, Jiangsu, Shandong, and Henan from16 July to 15 August.

### 2.4 Probability distribution of *S. frugiperda* entering Japan and Korea

Only trajectories of *S. frugiperda* that entered Japan and Korea were selected to make the distribution of an immigration probability, or invasion risk. The spatial frequency distribution of final endpoints of trajectories was calculated and displayed by using QGIS (Version 3.6, https://www.qgis.org/) and R (Version 3.5, https://www.r-project.org/), and this was achieved by the following procedures. Firstly, the Korean Peninsula and Japan were divided into several hexagonal cells, and the number of endpoints located in each hexagonal cell were counted in QGIS. Secondly, hexagonal cells in Japan (or in the Korean Peninsula) were divided equally into four groups based on the number of endpoints, (i) ≤ 25^th^ percentile, (ii) > 25^th^ percentile but ≤ median value, (iii) > medial value but ≥ 75^th^ percentile, and (iv) ≥ 75^th^ percentile. Finally, these results and the spatial distribution of trajectories’ origins were displayed by using R.

### 2.5 Wind pattern during migration season

To investigate wind pattern (wind speed and direction) in the last five years (2014–2018), we plotted average meteorological conditions at 850 hPa (approximately 1500 m above sea level). Long-term data were extracted and calculated from the daily NCEP/National Centre for Atmospheric Research (NCAR) Reanalysis data, with a spatial resolution of 2.5° by 2.5° global grids, and displayed using the Grid Analysis Display System (GrADS, Version 2.0.1, http://cola.gmu.edu/grads/).

## 3 RESULTS

To estimate the invasion risk of *S. frugiperda* into Korean Peninsula and Japan islands, 138780 forward trajectories were calculated. In total, 18082 (13.03%) trajectories reached one of these two regions (Fig. 2).

**Fig. 1:**
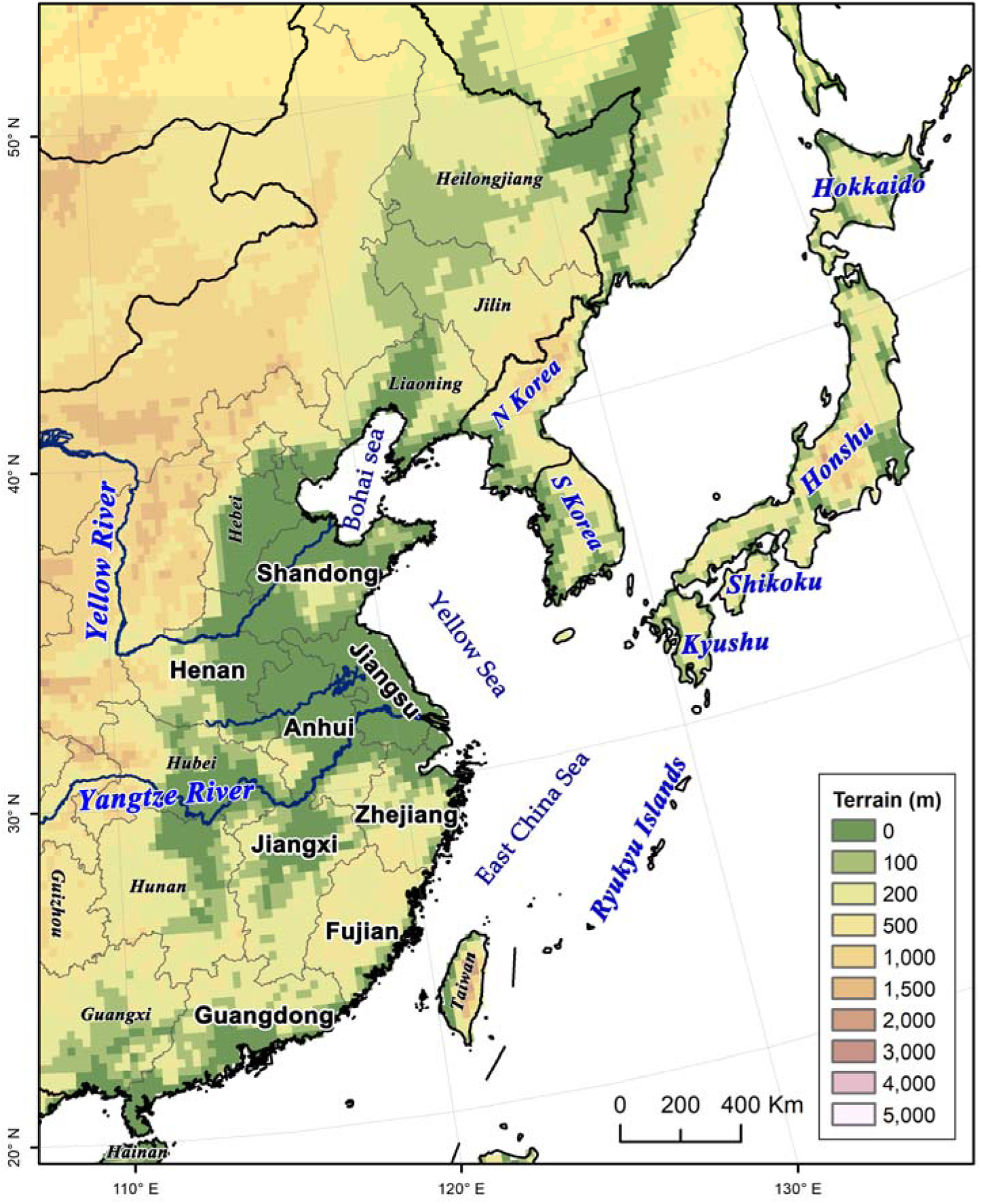
The map shows the topography of our study area in East Asia. Japan and the Korean Peninsula (South Korea and North Korea) were parts of the animal migration pathway in eastern Asia, as well as east China. *S. frugiperda* would migrate from east China into Japan and Korea by crossing sea, just like other migratory insects, such as rice planthoppers, rice leaf roller and oriental armyworm. Simulated trajectories of *S. frugiperda* start from Guangdong, Jiangxi, Fujian, Zhejiang, Anhui, Jiangsu, Henan, and Shandong provinces, which were shown in bold.

**Fig. 2:**
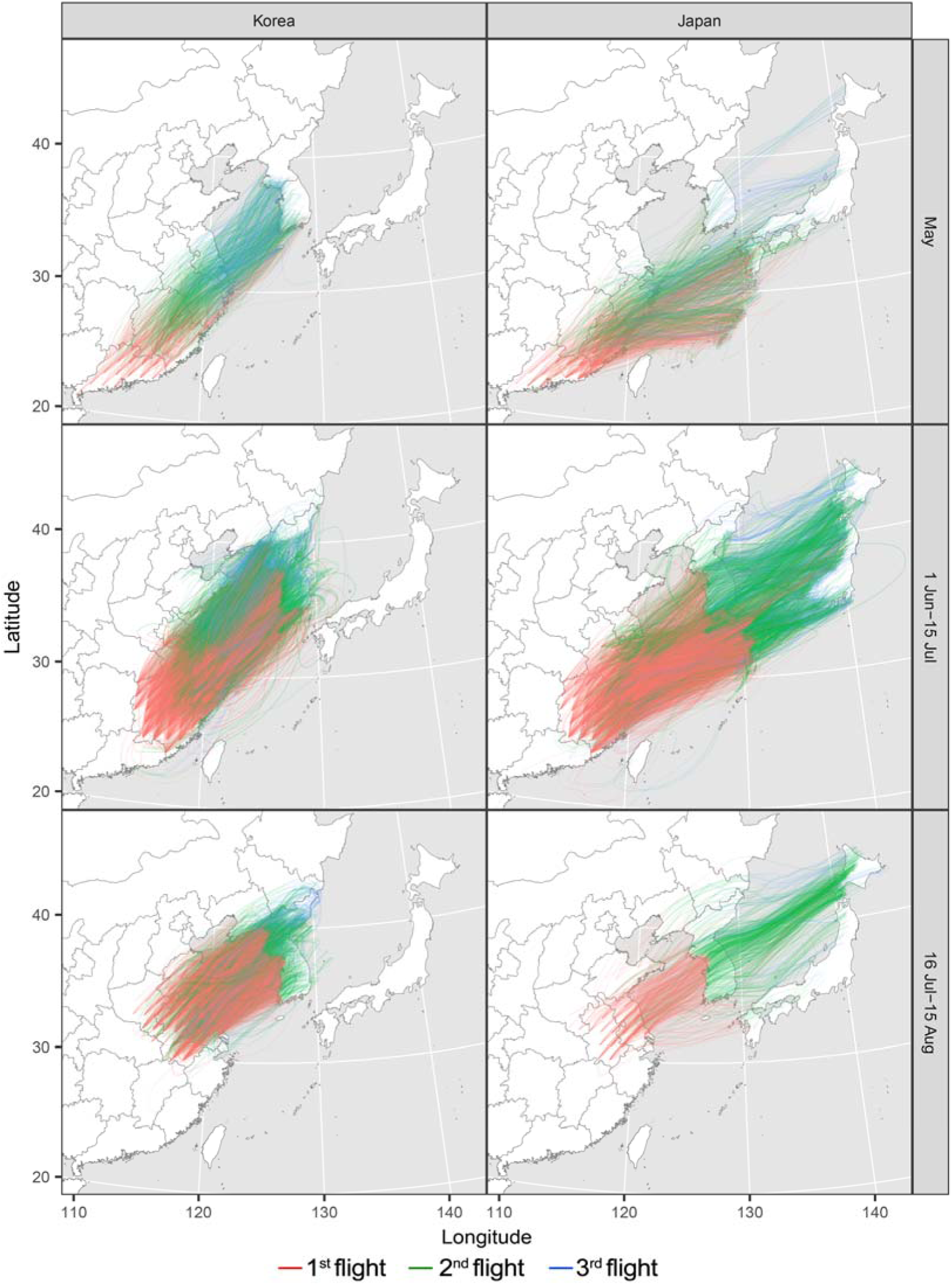
Forward trajectories of *S. frugiperda* reached the Korean Peninsula and Japan. The moth was assumed fly for 10 h over the land every night. When *S. frugiperda* comes over the sea area during each nocturnal flight, flight duration is extended until it reaches over the land but does not exceed 36 hours. *S. frugiperda* moth was assumed to take fly for three consecutive nights, but some overseas migrations were extended.

### 3.1 Immigration pattern on the Korean Peninsula

In May, 25110 trajectories (31 nights × 27 departure points × 6 initial heights × 5 years) were calculated and departed from Guangdong and Fujian. Among these, 698 (2.78%) trajectories arrived Korean Peninsula. Among these effective trajectories, their endpoints were located in the western and southern coastal areas of South Korea (Fig. 3). The straight distance between the departure and end points of each trajectory was 1631.76±10.88 km (mean value ± standard error, same after that). The flight duration to cover this distance was 30.81±0.34 h (Fig. 4).

**Fig. 3:**
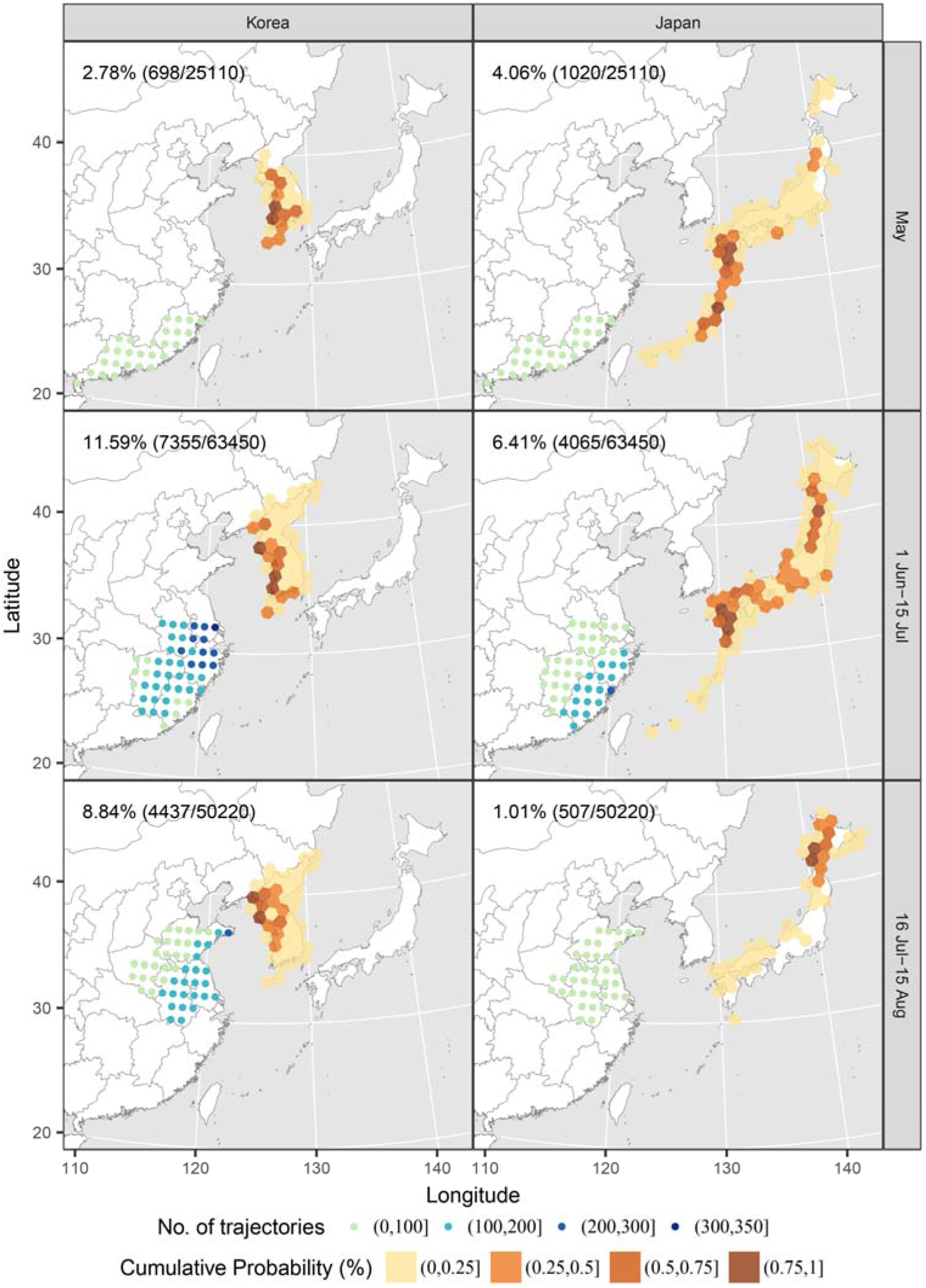
Distribution of start-points (dots in eastern China) and endpoints (hexagonal cells) of *S. frugiperda* forward migration trajectories reached Japan and Korea during every night of 1 May to 15 August of 2014–2018. The percentage of endpoints that successfully reached Japan or Korea is given with the number of all trajectories shown in parenthesis. Trajectory analyses were conducted over three consecutive nights normally. Here, only the final endpoint of each trajectory is shown. Each hexagonal and circular crossed every 100 × 100 km grid cell in the region.

**Fig. 4:**
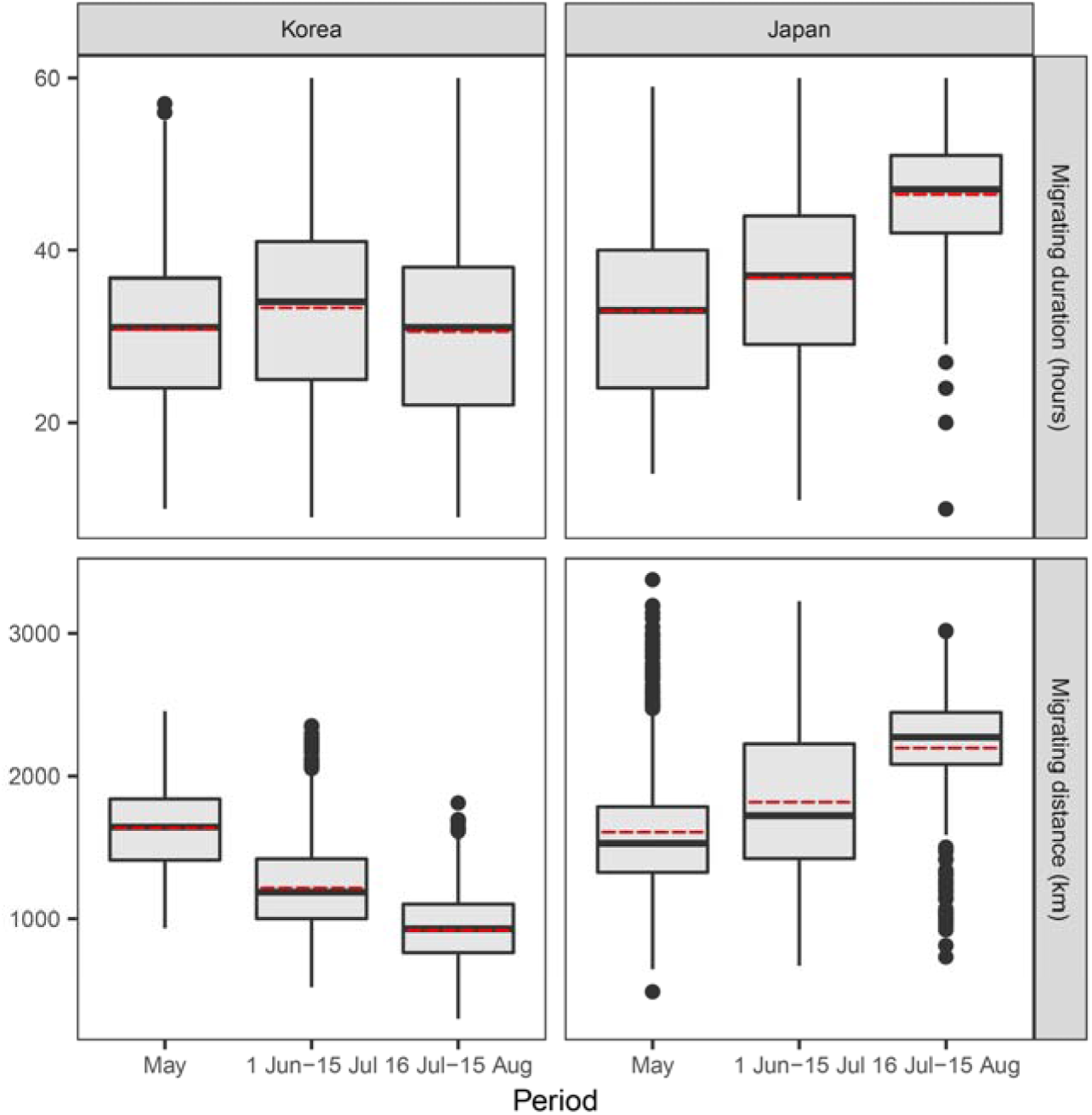
Flight duration and distance of *S. frugiperda* simulated trajectories reached Japan and Korea in different periods. The migrating distances mean straight distances between the start-points and the final endpoints for each trajectory. The solid black line represents median values, the red dashed line represents the mean, boxes represent the inter-quartile range (IQR), whiskers extend to observations within ±1.5 times the IQR, and dots represent outlier.

In the second period from June 1 to July 15, 63450 trajectories (45 nights × 47 departure points × 6 initial heights × 5 years) were calculated from Jiangxi, Zhejiang, and southern Anhui (south of 32°N), south Jiangsu (south of 32°N). Among these, 7355 (11.59%) arrived in the Korean Peninsula. Most of these effective trajectories originated from southern Jiangsu, southeastern Anhui, and northern Zhejiang (darker blue dots in Fig. 3). Their endpoints mostly located in the western coastal area of the peninsula (Fig. 3). The straight distance of these simulated trajectories was 1214.49±3.31 km, and the flight duration was 33.28±0.12 h (Fig. 4).

In the third period on 16 July to 15 August, 8.84% (4437/50220) trajectories originated from 54 departure points in Anhui, Jiangsu, Henan, and Shandong, reached the Korean Peninsula (Fig. 3). The endpoints of these effective trajectories distribute in the western part of North Korea and the northwestern part of South Korea. The main departure points distributed in eastern Shandong, Anhui, and Jiangsu (Fig. 3). The straight distance of these simulated trajectories was 922.80±3.68 km, and the flight duration was 30.50±0.15 h (Fig. 4).

In summary, the results predicted that *S. frugiperda* would invade into the Korean Peninsula, especially in the western coastal area. *S. frugiperda* in Anhui, Jiangsu and Shandong provinces would make main contributions to this invasion in the second (1 June–15 July) and third periods (16 July–15 August) (Fig. 3 & 4).

### 3.2 Immigration pattern on Japan

The percentages of trajectories from eastern China into Japan were 4.06% (1020/25110), 6.41% (4065/63450) and 1.01% (507/50220) on 1–31 May, 1 June– 15 July, and 16 July–15 August, respectively (Fig. 2 & 3). This result indicated it is quite possible that *S. frugiperda* would invade into Japan from 1 June through 15 July, and there was lower attainable in the other two periods. Kyushu, Shikoku, and southwestern Honshu and Shikoku would face a high risk of the invasion because most these trajectories ended over this region from 1 June to 15 July (Fig. 3). *S. frugiperda* invader into Japan would mostly come from Fujian and Zhejiang province (Fig. 2 & 3). Moreover, the Ryukyu Islands would be reached by *S. frugiperda* originated from Guangdong and Fujian provinces in May (Fig. 2 & 3). By contrast, quite lots of *S. frugiperda*’s trajectories reached Hokkaido by crossing Korean Peninsula in 16 July–15 August (Fig. 2 & 3).

During the three periods, 1–31 May, 1 June–15 July, and 16 July–15 August, the straight flight distances of the trajectories were 1602.77±13.67 km, 1818.32±7.95km, and 2193.99±18.05 km, respectively (Fig. 4). The flight duration to cover these distances were 32.91±0.32 h, 36.81±0.17 h, and 46.40±0.34 h, respectively (Fig. 4).

Take the results together, Japan also has a high risk of *S. frugiperda* invasion, especially in the second period, i.e., 1 June– 15 July. However, the possibility of the invasion into Japan was relatively lower when compared with the Korean Peninsula, because the overseas distance between eastern China and Japan were longer than that between eastern China and Korea (Fig. 3 & 4).

### 3.3 Wind pattern during migration season

The wind pattern on an isobaric level of 850 hPa (about 1500 m above sea level) during the migration season was investigated with a five-year dataset (2014–2018).

Southwest winds predominated in southern China and extended into Japan in May. Strong south-westerly air currents (≥ 4 m/s) flowed just in the Ryukyu Islands, and this indicated the lack of suitable winds to transport *S. frugiperda* into the mainland of Japan and the Korean Peninsula (Fig. 5a). Meanwhile, Guangdong and Fujian, the source area of *S. frugiperda*, are far from the mainland of Japan and the Korean Peninsula (Fig. 3). Thus, the frequency for *S. frugiperda* to reach the mainland of Japan and the Korean Peninsula was relatively lower in May, but the Ryukyu Islands can be reached by quite lots of *S. frugiperda* trajectories (Fig. 3 & 5a). In contrast, *S. frugiperda* in eastern China could spread very quickly in May due to strengthening southwesterly winds (Fig. 5a), and this scenario is exactly what happened until May of this year. After colonization in Yunnan, Hainan, Guangxi, and Guangdong, *S. frugiperda* appeared in Hunan, Hubei, Jiangxi, Fujian, and Zhejiang in May 2019 (NATESC, 2019c).

**Fig. 5:**
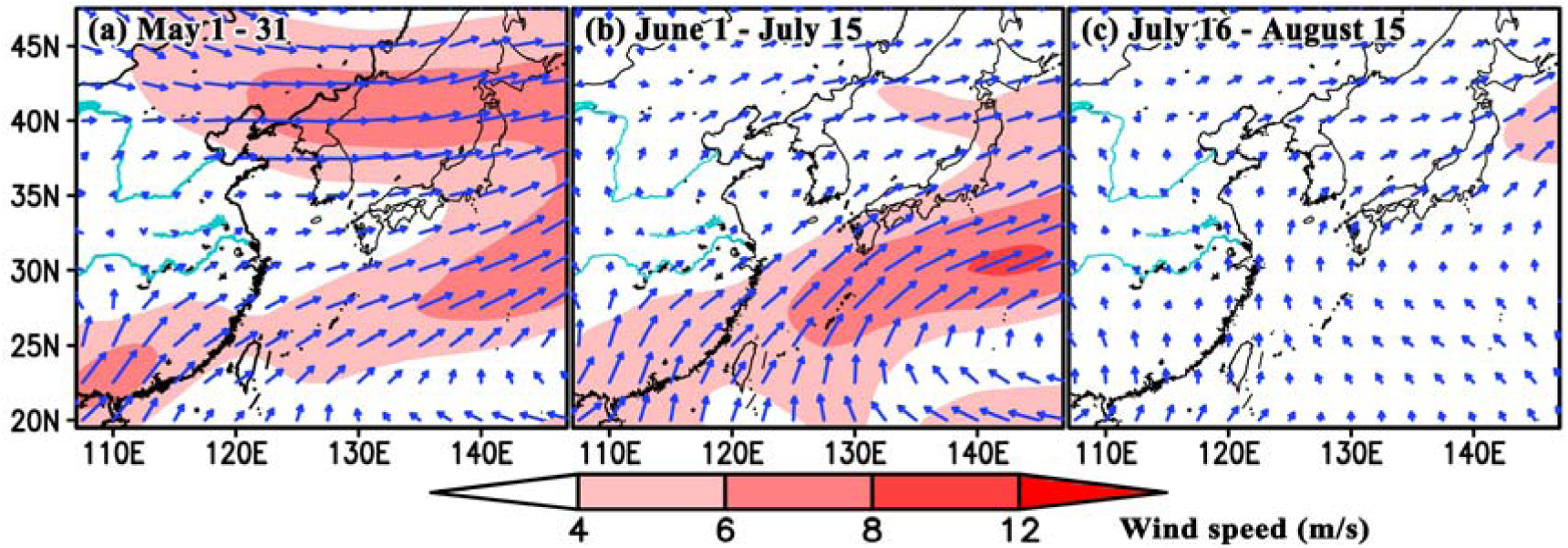
The five-year mean wind pattern at the level of 850 hPa (about 150 m above sea level) during migration periods of 2014–2018 over eastern Asia.

During 1 June–15 July, southwesterly winds strengthened and extended over East China Sea and Yellow Sea, and this would be helpful for *S. frugiperda* migrating from eastern China into Japan and Korean Peninsula (Fig. 5b). Although the mean wind speed over the Yellow Sea is quite lower (<4 m/s) (Fig. 5b), the distance between Jiangsu and Korean Peninsula is not very far (Fig. 3 & 4). As a result, larger proportion of simulated trajectories arrived in Japan, and Korean Peninsula than in other periods (Fig. 2 & 3).

The wind got weaken after 16 July. The wind direction changed to northward on the south of the Yellow River (Fig. 5c), and thus this was not suitable for *S. frugiperda* migrating into Japan. Eastward winds mostly appeared in northern China, Bohai Sea, and northern Yellow Sea (Fig. 5c), and this would help for *S. frugiperda*’s oversea migration into the Korean Peninsula. The wind speed was low (Fig. 5c), but the distance from Shandong and Jiangsu to the Korean Peninsula is relatively short (Fig. 4). It is the season that 8.84% of simulated trajectories reached the peninsula.

## 4 DISCUSSION

This study predicted the immigration risk of *S. frugiperda* into Japan and Korea by the trajectory analysis considering the flight behaviour of *S. frugiperda* based on the five-year meteorological data. The results showed that *S. frugiperda* would migrate from eastern China into Japan and Korea Peninsula shortly (Fig. 3).

Possibility that such overseas migrations will actually occur, or validity of the predicted risk in this study, depends mainly on the accuracy of assumptions made on the source location and the flight behaviour in Section 2. The rapid spreading of the moth in China started from southwestern Yunnan Province where the moths were firstly found in late January this year, and only within 3 months since then, which corresponds to the insect’s 3 generations, *S. frugiperda* was found in 13 provinces (NATESC, 2019a, b, c), including Guangdong, Fujian, Jiangxi and Zhejiang Provinces that were assumed as a source in this study. Because the moth’s spreading speed is very large, it is fair to predict that other assumed source provinces in Section 2.3 would be invaded by the rainy season and later summer.

Regarding the flight behaviour, trajectory analysis of the species in the previous study used such parameters as initial height of 500 m, flight duration of 12 h and 3-night flight for migration over the land of North America (Westbrook, Nagoshi, Meagher, Fleischer & Jairam, 2016). A previous trajectory analysis on the overseas migration of the common cutworm, *Spodoptera litura*, over East China Sea assumed flight height of 1000 and 1500 m, flight duration of 24 or 36 h (Tojo et al. 2013). This study assumed six initial heights from 500 to 1750 m, flight duration of 10 h over the land, up to 36 h in case of overseas migration, and 3-night flight as described in Section 2.3. Comparing with above these parameters each other, this study’s assumptions on the source and the flight behaviour seem reasonable. The invasion risk of this study, therefore, should be a fair prediction.

First possible arrivals in Japan could happen on southwestern small islands of Okinawa and Kagoshima Prefectures as well as western parts of Kyushu in May (Fig. 3b). Most likely, *S. frugiperda* would migrate from Fujian and Zhejiang into Kyushu, Shikoku and southern Honshu from 1 June – 15 July, the rainy season (Fig. 3d). During the summer period from mid-July, the invasion risk in northern Japan could relatively increase (Fig. 3f). As a whole, a wide area of Japan has an immigration risk.

First possible arrivals from Guangdong and Fujian could also occur in May in the western coastal areas of the Korean Peninsula (Fig. 3a). In the later periods of 1 June – 15 July and 16 July – 15 August, western Korean Peninsula would be continuously reached by *S. frugiperda* migrants from northern Zhejiang, Anhui, Jiangsu, and Shandong (Fig. 3c, e). According to the shift of the location of these possible sources from south to north, relatively higher risk also shifted from southern, middle to northern coastal areas of the peninsula (Fig. 3a, c, e).

These possible migrations could result in substantial damages and economic losses to agricultural production in Japan and Korea unless *S. frugiperda* population was effectively managed. Sugar cane, *Saccharum officinarum* is grown as a major agricultural product on the southwestern small islands. In Kyushu and mountainous areas in Honshu, livestock industry is active and maize, *Zea mays* L., for feed would a possible host plant for the immigrants. Maize is a major crop in Hokkaido, northern Japan, too. In northern part of the Korean Peninsula, maize is widely cultivated as one of important crops which would be severely damaged by summer migrants as predicted in Northeast China (Li et al., 2019). Sorghum (*Sorghum bicolor*) and hog millet (*Panicum miliaceum*) in northern Korea would be also attacked. If summer immigration would occur, plant protection officers in these areas should bear that possibility in mind. There are two types of biological strain of *S. frugiperda*, corn-strain and rice-strain (Pashley, Quisenberry & Jamjanya 1987). If the rice-strain to attack rice arrived in Japan and Korea, the almost all areas would be vulnerable to immigrants, because rice is cultivated everywhere in the nations. So far, Chinese population caught in Yunnan Province has been reported to be the corn-strain (Zhang et al., 2019).

Many species of insects can migrate into Japan and the Korea Peninsula by crossing over the sea, such as *S. litura, M. separata*, and rice planthoppers (Otuka et al., 2006; Otuka, 2013; Kisimoto & Sogawa, 1995; Hirai, 1995; Lee & Uhm, 1995; Tojo et al., 2013). *N. luges* and *S. furcifera* generally immigrate into Japan during the *Bai-u* rainy season from late June to early July (Kisimoto & Sogawa, 1995; Otuka et al., 2006). Kyushu and southern Honshu have the highest density of these two rice planthoppers (Kisimoto, 1976). The origin areas of these migrants was estimated by trajectory method to be southeastern China, such as Fujian, Zhejiang and Taiwan provinces (Otuka et al., 2006). In South Korea, *N. luges* also was reported to arrive in this same period of each year (Lee & Uhm, 1995; Zhu et al., 2000). In addition, *S. litura* is the same genus as *S. frugiperda*, and many male moths were caught by pheromone traps in Japan from June to middle July, and these moths were also thought to migrate from southern China (Tojo et al., 2013). Here, our study also suggested *S. frugiperda* would most likely carry out overseas migration into Japan and South Korea via similar pathway at similar time period (Fig. 2). The season of all those insect species migrating into Japan and South Korea at same time is that warm and humid southwesterly winds was quite frequent during the *Bai-u* season. This kind of wind always associated with the passage of depressions along the frontal zone, and the speed can up to 10 m/s or more (Kisimoto, 1976; Watanabe, Seino, Kitamura, Hirai, 1990; also see Fig. 4).

Conversely, the migration pathway and timing of *L. striatellus* and *M. separata* are different with those aforementioned species. A single-flight migration route of *M. separata* invaded into Korea and northern Japan mainly in late May – early June is suggested in previous studies (Haria, 1995; Lee & Uhm, 1995). The moths might originate in eastern China at 30–36°N (probably from Jiangsu, Anhui, Henan and Shandong) (Jiang et al., 2011), crossing over the northern Yellow Sea, passing beyond the Korean Peninsula at 37–38°N, and arriving in northern Japan at 39–p45°N (Hirai, 1995; Lee & Uhm, 1995). Similarly, *L. striatellus* from Jiangsu migrating into western Japan and western South Korea was also reported by crossing over the Yellow Sea in late May and early June (Otuka, 2013). Immigrations of both these species occurs most frequently on strong southwesterly or westerly winds, also associated with the passage of a low pressure system (Haria, 1995; Lee & Uhm, 1995; Otuka, 2013). *S. frugiperda* would likely have similar migration route but at different timing, in 16 July–15 August. This is presumably because *S. frugiperda* appeared in Jiangsu, Anhui and Shandong later than the former two species. *L. striatellus* can survive cold winter in most area of eastern China, even in the Northeast Plain (Sun et al., 2015), and the population in Jiangsu emigrates when wheat matures and was harvested in late May (Otuka, 2013). *M. separata* overwinters in a region south of the Yangtze River (< 33 ºN) in China (Sun, 1990), and reach the region of 30–34°N in March and April (Chen, 1995; Jiang et al., 2011). However, the year-round distribution of *S. frugiperda*, which does not possess a capability to enter diapause (Westbrook et al., 2016), in East Asia is restricted to relatively warm and moist regions found on the Indochina Peninsula and in southern China (to the south of the Tropic of Cancer) (Early et al., 2018), and thus it would arrive in Jiangsu and Shandong later.

In conclusion, Japan and Korea face a high risk of invasion of *S. frugiperda* mainly in 1 June–15 July, the *Bai-u* rainy season. This migration pattern was also used by many other migratory insects. *S. frugiperda* in eastern China actually spread very quickly in May (NASTEC 2019c), and this coincided precisely with the predictive range from our previous study with the same approach (Li et al., 2019). Therefore, the continuing spread through China to Japan and Korea seems inevitable. Additional studies on its migration patterns, flight behaviour, ecology, and pest management based on monitoring data in East Asia are urgently required.

## ACKNOWLEDGMENTS

This work was supported though grants to G.H. by the National Natural Science Foundation of China (31822043) and the Natural Science Foundation of Jiangsu Province (BK20170026). B.G.’s visiting scholarship to the University of Exeter was funded by the China Scholarship Council and the Jiangsu Graduate Research and Innovation Projects.

## DECLARATION OF INTERESTS

The authors declare that they have no competing interests.

## AUTHOR CONTRIBUTION

GH, AO and GL conceived the research plan. JM, YW and MW conducted the trajectory analysis. JM, BG, GH, AO and GL wrote the manuscript. GH secured funding. All authors read and approved the manuscript.

## REFERENCES

Casmuze, A., Juárez, M. L., Socías, M. G., Murúa, M. G., Prieto, S., Medina, S., Willink, E. & Gastaminza, G. (2010). Review of the host plants of fall armyworm, *Spodoptera frugiperda* (Lepidoptera: Noctuidae). Rev. Soc. Entomol. Argent. 69, 209–231.

Chen, R.L., Sun, Y. J., Wang, S. Y., Zhai, B. P., Bao, X. Y. (1995) Migration of the oriental armyworm *Mythimna separata* in East Asia in relation to weather and climate. I. Northeastern China, In VA Drake & AG Gatehouse (Eds.), Insect Migration: Tracking Resources through Space and Time (pp. 93–104). Cambridge University Press, Cambridge, UK.

Cock, M. J. W., Beseh, P. K., Buddie, A. G., Giovanni Cafá. & Crozier, J. (2017). Molecular methods to detect *Spodoptera frugiperda* in Ghana, and implications for monitoring the spread of invasive species in developing countries. Scientific Reports, 7(1). https://doi.org/10.1038/s41598-017-04238-y

Early, R., Gonzalez-Moreno, P., Murphy, S. T. & Day, R. (2018). Forecasting the global extent of invasion of the cereal pest *Spodoptera frugiperda*, the fall armyworm. NeoBiota, 40, 25–50.

FAO. (2018–06–28) [2019-05-07]. Fall armyworm keeps spreading and becomes more destructive [EB/OL]. https://doi.org/www.fao.org/news/story/en/item/1142085/icode/

Goergen, G., Kumar, P. L., Sankung, S. B., Togola, A., Tamò, Manuele. & Luthe, D. S. (2016). First report of outbreaks of the fall armyworm *Spodoptera frugiperda* (J E Smith) (Lepidoptera, Noctuidae), a new alien invasive pest in west and central Africa. PLoS ONE, 11(10). https://doi.org/10.1371/journal.pone.0165632

Guo, J. F., Zhao, J. Z., He, K. L., Zhang, F. & Wang, Z. Y. (2018). Potential invasion of the crop-devastating insect pest fall armyworm *Spodoptera frugiperda* to China. Plant Protection, 44(6), 1–10.

Hirai, K. (1995). Migration of the Oriental Armyworm *Mythimna separata* in East Asia in relation to weather and climate. III. Japan, In VA Drake & AG Gatehouse (Eds.), Insect Migration: Tracking Resources through Space and Time (pp. 117–129). Cambridge University Press, Cambridge, UK.

Hogg, D. B., Pitre, H. N. & Anderson, R. E. (1982). Assessment of early-season phenology of the fall armyworm (Lepidoptera: Noctuidae) in mississippi [*Spodoptera frugiperda*]. Environmental Entomology, 11(3), 705–710. https://doi.org/10.1093/ee/11.3.705

Jaeyoul, U., Donghyuck, L., Sangkye, L., Uhm, J. Y., Lee, D. H. & Lee, S. K. (1995). Development of fungicide spray program for the apples to be exported to the united states of america. Korean Journal of Plant Pathology, págs. 83–104.

Jiang X., Luo, I., Zhang, L., Sappington, T. W. & Hu, Y. (2011). Regulation of migration in *Mythimna separata* (Walker) in China: A review integrating environmental, physiological, hormonal, genetic, and molecular factors. Environmental Entomology, 40(3), 516–533. https://doi.org/dx.doi.org/10.1603/EN10199.

Jiang, Y. Y., Liu, J. & Zhu, X. M. (2019). Analysis on the occurrence dynamics of invasion and future trend of fall armyworm *Spodoptera frugiperda* in China. China Plant Protection, 39(2), 33-35.

Johnson, S. J. (1987). Migration and the life history strategy of the fall armyworm, *Spodoptera frugiperda* in the western hemisphere. International Journal of Tropical Insect Science, 8 (4-5-6), 543–549. https://doi.org/10.1017/S1742758400022591

Kisimoto, R. & Sogawa, K. (1995). Migration of the Brown Planthopper *Nilaparvata lugens* and the White-backed Planthopper *Sogatella furcifera* in East Asia: the role of weather and climate, In VA Drake & AG Gatehouse (Eds.), Insect Migration: Tracking Resources through Space and Time (pp. 61–91). Cambridge University Press, Cambridge, UK.

Kisimoto, R. (1976). Synoptic weather conditions inducing long-distance immigration of planthoppers, *Sogatella furcifera Horvath* and *Nilaparvata lugens Stål*. Ecological Entomology, 1, 95–109. https://doi.org/10.1111/j.1365-2311.1976.tb01210.x

Lee, J. H. & Uhm, K. B. (1995). Migration of the Oriental Armyworm Mythimna separata in East Asia in relation to weather and climate. II. Korea., In VA Drake & AG Gatehouse (Eds.), Insect Migration: Tracking Resources through Space and Time (pp. 105–116). Cambridge University Press, Cambridge, UK.

Li, X. J., Wu, M. F., Ma, J., Gao, B. Y., Wu, Q. L., Chen, A. D., Liu, J., Jiang, Y. Y., Zhai, B. P., Early, R., Chapman, J. W. & Hu, G. (2019-05-02) [2019-05-07]. Prediction of migratory routes of the invasive fall armyworm in eastern China using a trajectory analytical approach [J/OL]. bioRxiv, https://www.biorxiv.org/content/10.1101/625632v1.

Luginbill, P. (1928). The fall armyworm. USDA Technology Bulletin, 34, 91.

Nagoshi RN, Meagher RL, Fleischer S. (2009) Texas is the overwintering source of fall armyworm in central Pennsylvania: implications for migration into the northeastern United States. Environmental Entomology, 38 (6), 1546–1554.

Nagoshi, R. N., Goergen, G., Tounou, K. A., Agboka, K., Koffi, D. & Meagher, R. L. (2018). Analysis of strain distribution, migratory potential, and invasion history of fall armyworm populations in northern sub-Saharan Africa. Sci Rep, 8(1), 3710. https://doi.org/10.1038/s41598-018-21954-1

National Agricultural Technology Extension Service Center (NATESC), (2019a) Major pest *Spodoptera frugiperda* have invaded in Yunnan, and all areas should immediately strengthen investigation and monitoring. Plant pathogen and pest information. 2019-1-18

National Agricultural Technology Extension Service Center (NATESC), (2019b) Recent reports of fall armyworm in China and neighbouring countries. Plant pathogen and pest information. 2019-4-4.

National Agricultural Technology Extension Service Center (NATESC), (2019c) Fall armyworm harms spring corn in 13 provinces in China. Plant pathogen and pest information. 2019-5-14.

Otuka, A. (2013). Migration of rice planthoppers and their vectored re-emerging and novel rice viruses in East Asia. Frontiers in Microbiology, 4, 309. https://doi.org/10.3389/fmicb.2013.00309

Otuka, A., Watanabe, T., Suzuki, Y., Matsumura, M., Furuno, A., Chino, M., Kondo, T. & Kamimuro, T. (2006). A migration analysis of *Sogatella furcifera* (Horváth) (Homoptera: Delphacidae) using hourly catches and a three-dimensional simulation model. AgriculturalandForestEntomology, 8(1), 35–47. https://doi.org/10.1111/j.1461-9555.2006.00284.x

Pashley, D. P., Quisenberry, S. S., Jamjanya, T. (1987). Impact of fall armyworm (Lepidoptera: Noctuidae) host strains on the evaluation of bermudagrass resistance. J Econ. Entomol 80, 1127–1130.

Qi Guo-Jun., Li-Hua, L. V., Ri-Qing, L., Jin-Hong, X. & Wei-Qun, Z. (2013). Tracking the source regions of cnaphalocrocis medinalis in the rice growing region of northern guangdong province. Chinese Journal of Applied Entomology, 50(3), 601–607

Rwomushana, I., Bateman, M., Beale, T., Beseh, P., Cameron, K., Chiluba, M., Clottey, V., Davis, T., Early, R., Godwin, J., Gonzalez-Moreno, P., Kansiime, M., Kenis, M., Makale, F., Mugambi, I., Murphy, S., Nunda, W., Phiri, N., Pratt, C. & Tambo, J. (2018). Fall armyworm: impacts and implications for Africa Evidence Note Update, October 2018. Report to DFID. Wallingford, UK: CAB International.

Sharanabasappa Kalleshwaraswamy, C. M., Asokan, R., Mahadeva, S. H. M., Maruthi, M. S., Pavithra, H.B., Hegde, K., Navi, S., Prabhu, S.T. & Goergen, G. (2018). First report of the fall armyworm, *Spodoptera frugiperda* (J E Smith) (Lepidoptera: Noctuidae), an alien invasive pest on maize in India. Pest Management: in Horticultural Ecosystems, 24(1), 23–29.

Sparks, A. N. (1979). A review of the biology of the fall armyworm. The Florida Entomologist, 62(2), 82–87. https://doi.org/10.2307/3494083

Stokstad. & Erik. (2017). New crop pest takes africa at lightning speed. Science, 356(6337), 473–474. https://doi.org/10.1126/science.356.6337.473

Sun JR. Field surveys on the overwintering of oriental armyworm migration. In Physiology and Ecology of Oriental Armyworm, ed. by Lin CS, Chen RL, Shu XY, Hu BH and Cai XM, *Peking University Press*, pp, 167–172 (1990).

Sun, J. T., Wang, M. M., Zhang, Y.K., Chapuis, M. P., Jiang, X. Y., Hu, G., Yan, X. M., Ge, C., Xue, X. F., Hong, X. Y. (2015). Evidence for high dispersal ability and mito-nuclear discordance in the small brown planthopper, Laodelphax striatellus. Scientific Reports, 5:8045.

Tojo, S., Ryuda, M., Fukuda, T., Matsunaga, T., Choi D.-R. & Otuka, A. (2013). Overseas migration of the common cutworm, *Spodoptera litura* (Lepidoptera: Noctuidae), from May to mid-July in east Asia. Applied Entomology and Zoology, 48(2), 131–140. https://doi.org/10.1007/s13355-013-0162-x

Wang, F. Y., Yang, F., Lu, M. H., Luo, S. Y., Zhai, B. P., Lim, K. S., McInerney, C. E. & Hu, G. (2017). Determining the migration duration of rice leaf folder (*Cnaphalocrocis medinalis (Guenée)*) moths using a trajectory analytical approach. Scientific Reports, 7: 39853. https://doi.org/10.1038/srep39853

Watanabe, T., Seino, H., Kitamura, C., Hirai, Y. (1990). A computer program, LLJET, utilizing an 850 mb weather chart to forecast long-distance rice planthopper migration. Bulletin of the Kyushu National Agricultural Experiment Station, 26, 233–260.

Westbrook & J. K. (2007). Noctuid migration in texas within the nocturnal aeroecological boundary layer. Integrative and Comparative Biology, 48(1), 99–106. https://doi.org/10.1093/icb/icn040

Westbrook, J. K., Nagoshi, R. N., Meagher, R. L., Fleischer, S. J. & Jairam, S. (2016). Modeling seasonal migration of fall armyworm moths. International Journal of Biometeorology, 60(2), 255–267. https://doi.org/10.1007/s00484-015-1022-x

Wolf, W. W., Westbrook, J. K., Raulston, J. R. & Lingren, P. D. (1995). Radar observations of orientation of noctuids migrating from corn fields in the lower rio grande valley. Southwestern Entomologist Supplement, 20(3),45–61. https://doi.org/10.1111/j.1365-3032.1995.tb00012.x

Wolf, W. W., Westbrook, J. K., Raulston, J., Pair, S. D., Hobbs, S. E. & Riley, J. R., et al. (1990). Recent airborne radar observations of migrant pests in the united states [and discussion]. Philosophical Transactions of the Royal Society B: Biological Sciences, 328(1251), 619–630. https://doi.org/10.1098/rstb.1990.0132

Wu, Q. L., Jiang, Y. Y. & Wu, K. M. (2019). Analysis of migration routes of fall armyworm, *Spodoptera frugiperda* (J. E. Smith) from Myanmar to China. Plant Protection, 45(2): 1–9.

Zhang, L., Jin, M. H., Zhang, D. D., Jinag, Y. Y., Liu, J., Wu, K. M. & Xiao, Y. T. (2019) Molecular identification of invasive fall armyworm *Spodoptera frugiperda* in Yunnan Province. Plant Protection, 45: 19–24.

Zhu, M., Song, Y. H., Uhm, K. B., Turner, R. W., Lee, J. H. & Roderick, G. K. (2000) Simulation of the long range migration of Brown Planthopper, *Nilaparvata lugens* (Stål), by using boundary layer atmospheric model and the geographic information system. J. Asia-Pacific Entomol. 3: 25–32.

